# Optogenetic stimulation of the VTA modulates a frequency-specific gain of thalamocortical inputs in infragranular layers of the auditory cortex

**DOI:** 10.1101/669168

**Authors:** Michael G. K. Brunk, Katrina E. Deane, Martin Kisse, Matthias Deliano, Silvia Vieweg, Frank W. Ohl, Michael T. Lippert, Max F. K. Happel

## Abstract

**Background:** Reward associations during auditory learning induce cortical plasticity in the primary auditory cortex. A prominent source of such influence is the ventral tegmental area (VTA), which conveys a dopaminergic teaching signal to the primary auditory cortex. It is currently unknown, however, how the VTA circuitry thereby influences cortical frequency information processing and spectral integration. In this study, we therefore investigated the temporal effects of direct optogenetic stimulation of the VTA onto spectral integration in the auditory cortex on a synaptic circuit level by current-source-density analysis in anesthetized Mongolian gerbils.

**Results:** While auditory lemniscal input predominantly terminates in the granular input layers III/IV, we found that VTA-mediated modulation of spectral processing is relayed by a different circuit, namely enhanced thalamic inputs to the infragranular layers Vb/VIa. Activation of this circuit yields a frequency-specific gain amplification of local sensory input and enhances corticocortical information transfer, especially in supragranular layers I/II. This effects further persisted over more than 30 minutes after VTA stimulation.

**Conclusions:** Altogether, we demonstrate that the VTA exhibits a long-lasting influence on sensory cortical processing via infragranular layers transcending the signaling of a mere reward-prediction error. Our findings thereby demonstrate a cellular and circuit substrate for the influence of reinforcement-evaluating brain systems on sensory processing in the auditory cortex.

## Background

The sensory cortex receives both bottom-up input relaying stimulus information from the sensory epithelia and top-down input from, for example, reinforcement-evaluating brain structures [1]. Among the latter, the ventral tegmental area (VTA) is a key structure associated with the coding of reward, reward prediction, and reward prediction error [2]. Especially in the framework of reward prediction error coding, projections of dopamine neurons in the VTA to the striatum and prefrontal cortex have been investigated in great detail [3]. In contrast, the anatomy of projections from the VTA towards sensory cortices [4–6], and consequently its direct impact on cortical processing, has remained rather elusive.

Dopamine released in the sensory cortex may complement bottom-up stimulus processing with a behaviorally relevant representation of stimulus value and salience to support adaptive behavior [2, 7, 8]. Consistently, for the case of the auditory cortex (ACx), intracortical dopamine levels [9] and their experimental manipulation [10, 11] were shown to affect behavioral measures in auditory learning scenarios. Thus, dopamine appears to be involved in several general behaviorally relevant functions, including auditory perceptual decision making, prediction, and learning, which are increasingly realized to be supported by the ACx [12]. In agreement, we have previously shown that the pharmacological stimulation of D1/D5 receptors influences sensory processing at the level of both local and wide-spread circuits in auditory cortex [13].

In order to determine the contribution of VTA projections to these cortical effects of dopamine, we optogenetically stimulated the projection neurons of the VTA in adult male Mongolian gerbils (*Meriones unguiculatus*) and measured the layer-specific processing in the auditory cortex by tone-evoked current-source density (CSD) analysis. We demonstrate that VTA stimulation effectuated a sensory gain amplification via thalamocortical inputs in the deep layers Vb/VIa, rather than via recurrent excitation in layer III/IV [14, 15]. Our results demonstrate for the first time a functional diversification of the anatomically distinct thalamocortical input systems in the sensory cortex. Reward-modulated sensory input in deep layer neurons therefore might provide a cellular substrate for integrating sensory and task-related information in the service of sensory-based decision-making and reinforcement learning.

## Results

### Optogenetic stimulation of the VTA evokes self-stimulation behavior

To selectively target the excitatory projection neurons of the VTA, we expressed the red-shifted opsin C1V1 under the CamKIIα-promotor [16] by a stereotactically guided microinjection (Fig. 1 A; 700 nl; AAV-CamKIIα-C1V1(E162T)-p2A-eYFP, 3e12 particles per ml, UNC Vector Core). Opsin expression in the VTA overlapped significantly with tyrosine hydroxylase (TH)-immunostaining, selective for dopaminergic cell populations. The connectivity of the transduced neurons with the auditory cortex was confirmed by histology (Fig. 1 A,B) in agreement with previous reports [4, 17]. After viral transduction, we waited 2-3 weeks for sufficient transgene expression before subjects were trained over the course of 10 days in an optogenetic intracranial self-stimulation task (Fig. 1 C; Additional file 1; 10 pulses, 25 Hz, 473 nm laser-light per nose-poke; 10 mW power at fiber implant) [18]. Animals that showed more than 50 presses/minute were chosen as the C1V1 group for electrophysiological experiments (Fig. 1 G) [19, 20]. Anatomical validation of the fiber position relative to Bregma (AP:−3.57 mm, ML: 584.14 μm; cf. Fig. 1 D and E), as well as the fiber distance towards the area of opsin expression (Fig. 1 F, 146.8 ± 38.9 μm), was performed in a subset of animals (n=7) in the C1V1 group (see Materials and Methods). In contrast to the opsin group, animals of the YFP-control group showed no self-stimulation behavior (Fig. 1 G).

**Figure 1 -.**
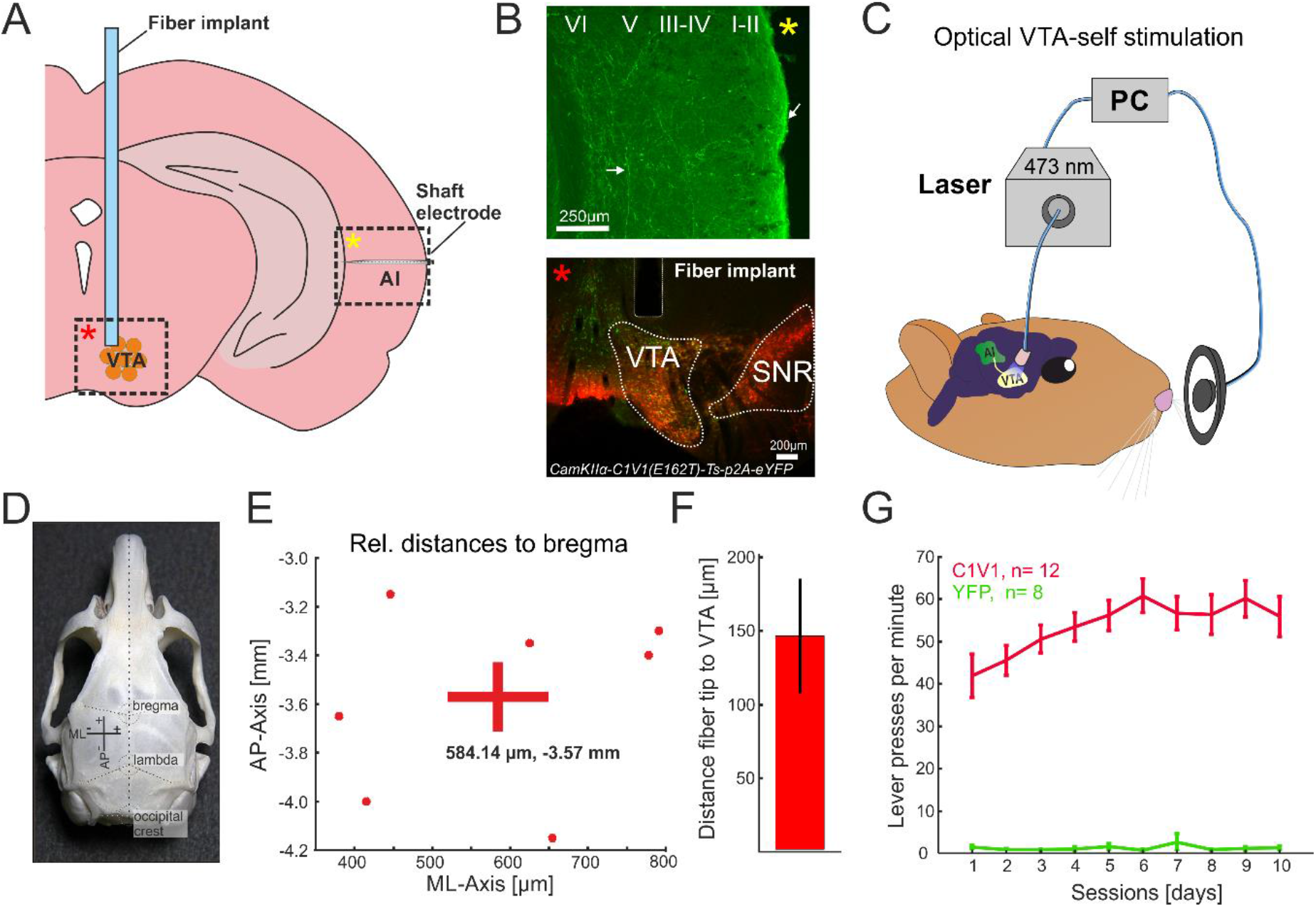
Anatomical validation of fiber position relative to target side and optogenetic intracranial self-stimulation. (A) Schematic overview of fiber implant position for optogenetic stimulation of the VTA, and the ipsilateral AI recording side. Dashed boxes indicate representative regions of interest shown in B) identified by yellow and red asterisks. (B) Top, YFP expression in the temporal cortex covering the region of AI after virus transduction of the VTA indicates fiber terminals from the VTA targeting mainly upper layers and collaterals in deeper layers (white arrows). YFP expression was detectable in multiple regions of the neocortex (not shown). Bottom, co-fluorescence of YFP (AAV-CamKIIα-C1V1 (E162T)-p2A-eYFP) and TH-immunostaining (Alexa Fluor 546). Area of co-fluorescence is indicative of overlapping neuronal populations expressing the virus and TH mainly found in the VTA, and not the substantia nigra [18]. Scale bars indicate 250 and 200 μm, respectively. (C) Schematic representation of optogenetic intracranial self-stimulation: While nose-poking the lever, animals induce a light flash towards the VTA resulting in extensive self-stimulating behavior [19]. (D) Gerbil skull with prominent landmarks (bregma, lambda and occipital crest) for anatomical reference [modified from 69]. For experiments, bregma has been used for reference of medio-lateral (ML) and anterior-posterior (AP) position of the VTA. (E) Estimated actual AP/ML coordinates with mean coordinates (AP:−3.57 mm; ML: 584.14 μm) relative to bregma of the C1V1group (n=7) as estimated with the gerbil brain atlas [69]. (F) Fiber distance (mean ± SEM) towards area of co-fluorescence (146.8 ± 38.9 μm; n=7). Compare with bottom picture of B). (G) Averaged lever pressing rates (lever presses/ min) of the C1V1 (n=12) and YFP (n=8) groups over 10 days. A representative video of a C1V1 animal is accessible as Movie (See Additional file 1).

### Effect of VTA-activation on tone-evoked columnar activity in the auditory cortex

After evaluation of self-stimulation behavior in the C1V1 and YFP group, we performed acute laminar local field potential electrophysiological recordings from their auditory cortices under ketamine-xylazine anesthesia. This approach allows the investigation of circuitry effects by the stimulation of the VTA on sensory cortical processing. Additionally, we also recorded from a control group of naïve animals, which underwent no implantation or laser stimulation, to determine which effects were due to opsin activation or laser light. Pure tones were presented in iso-octave steps in order to cover the main tonotopic range of the gerbil (Stimulation frequencies ranged from 0.125 kHz – 32 kHz, duration: 200 ms, ISI 0.6-0.8s, 50 rep., 65 dB SPL). After insertion of the recording electrode (NeuroNexus A1×32-50-413), sets of 50 repetitions per stimulus (~7.5 minutes) were recorded over 45-75 min to guarantee stable recordings (Fig. 2 A; see Materials and Methods). We then paired the presentation of each pure tone with laser-stimulation of the VTA (473 nm; 25 Hz; 10 ms pulse length, 200 ms duration). Thereby, the VTA was stimulated 400 times within 7.5 minutes of the combined laser/tone presentation (Fig. 2). After this paired measurement, consecutive tone-only measurements (post) were performed over the course of 60 minutes. This procedure allowed us to observe phasic and long-lasting effects of optogenetic VTA stimulation on auditory cortex tone-evoked activity.

**Figure 2 -.**
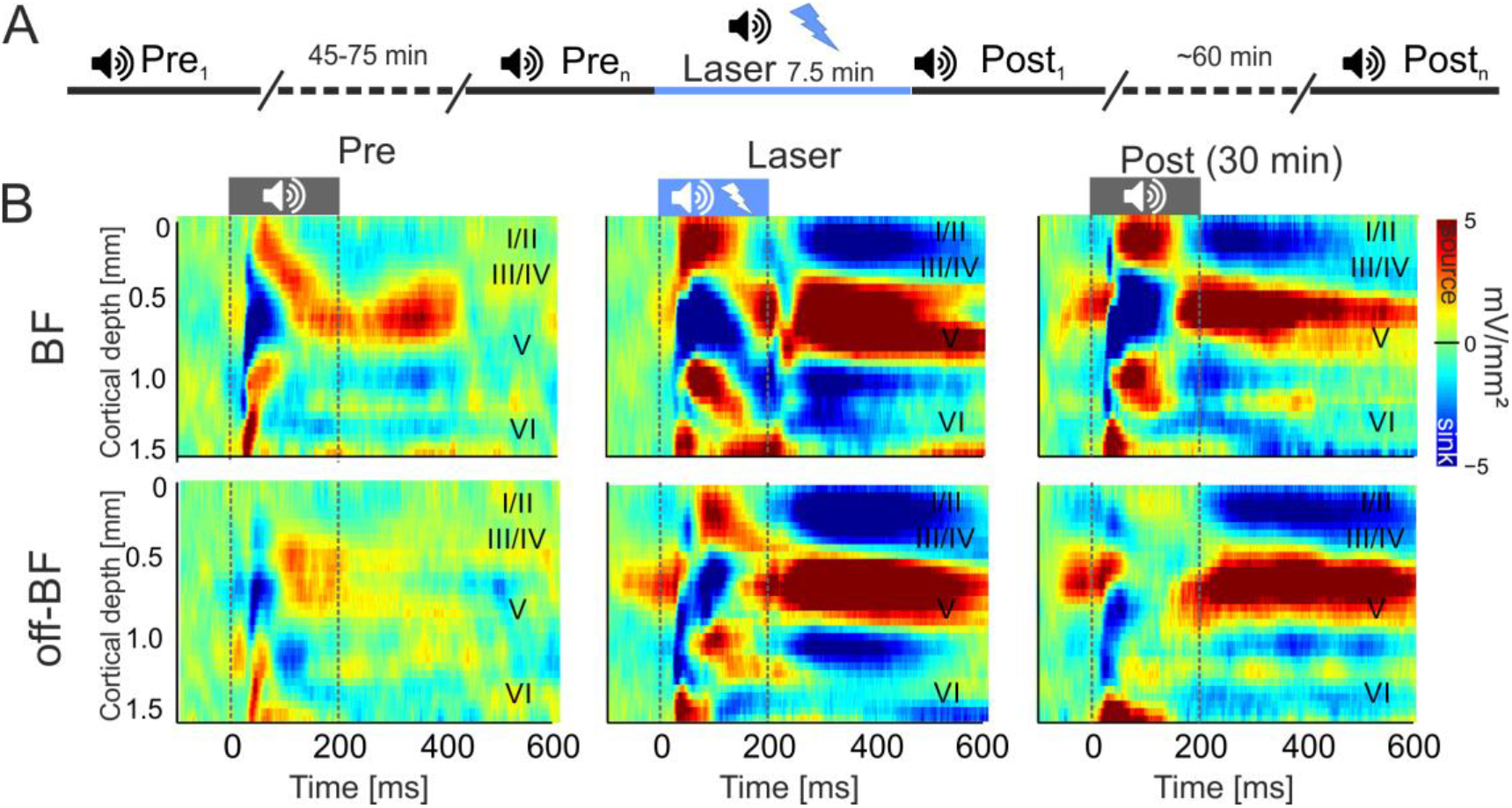
Electrophysiological measurements and effects of VTA stimulation on tone-evoked CSD profiles in A1. (A) Schematic representation of the timeline of measurements. Pre baseline measurements (Tone stimulation: 0.125-32 kHz, tone duration: 200 ms, ISI: 0.6-0.8, 50 pseudorandomized repetitions, 65 dB SPL, 7.5 min) were performed until CSD patterns had stabilized (45-75 min). A single measurement of combined tone and laser stimulation (25 Hz, 473 nm, 10 mW) was carried out and followed by a series of post-measurements (up to 60 min). (B) Representative example of a CSD profile after BF- (top) and off-BF (bottom) stimulation before, during and 30 minutes after the combined tone/VTA-laser measurement from an animal of the C1V1 group. Before VTA stimulation, BF-evoked CSD profiles revealed a canonical feedforward pattern of stimulus-evoked current flow starting with initial sink components in cortical layers III/IV and Vb/VIa. Subsequent sink activity was found in layers Va, I/II and VIb. Activity after off-BF stimulation showed a less pronounced columnar activation in the pre-condition. During paired tone and VTA stimulation, both stimulation frequencies evoked stronger and prolonged translaminar current flow, which was still enhanced 30 minutes later. Cortical layers are indicated. Dashed vertical lines indicate tone onset and offset.

Fig. 2 B displays a representative example of pure-tone evoked CSD-profiles after stimulation with the best frequency (BF, top) and an off-BF (bottom, 2 octaves apart) before (left), during (center) and 30 minutes after (right) VTA-stimulation in a C1V1-transduced animal. Before VTA stimulation, tone-evoked CSD profiles show the canonical feed-forward pattern of early current sink activity in layers III/IV and Vb/VIa (Fig. 2 B, left) [15, 21, 22]. Subsequent later supra- and infragranular sink activities can be related to intracortical synaptic circuits in layers I/II, Va and VIb (see semi-automatic detection of sink activity in Materials and Methods and Fig. 2 B and Additional file 2). Responses to frequencies apart from the BF generally evoke less prominent tone-evoked activity across all cortical layers. When pure tone stimulation is paired with VTA stimulation, we observed prominent changes of amplitudes, but not spatiotemporal flow of synaptic activity (Fig. 2 B, center). This increase of tone-evoked cortical processing was long-lasting and persisted over approximately 30 min after the cessation of laser stimulation (Fig. 2 B, right).

### Laminar specificity of transient and long-lasting effects of VTA stimulation

In order to relate the observed changes of in current flow to distinct cortical layers, we separately analyzed CSD-traces from respective cortical layers (I/II, III/IV, Va, Vb/VIa and VIb; see Materials and Methods). Prominent sink components of cortical layers III/IV, Va, and Vb/VIa were related to early time windows of 0-50 ms after tone onset, while cortical layers I/II and VIb felt into later time windows of >50 ms determined by sink onset (cf. Additional file 2).

Described effects during and after VTA stimulation might impact on sensory processing in a less stimulus-locked manner and influence trial-by-trial signal-to-noise ratio [23]. In order to capture such variability, we have utilized a single-trial analysis using LME models [24] on peak amplitudes of each layer as a function of time before and after VTA stimulation separately for BF, near- (± 1-2 octaves) and non-BF (± 3-4 octaves) stimulation for all groups (Fig. 3). Within each frequency bin, data was normalized to pre-measurements in order to characterize relative changes over time and compared both groups with laser treatment (C1V1, YFP) against the control group. Within the control group, we observed a variation of tone-evoked responses of up to ± 10% over the entire recording time. Therefore, to obtain main effects of the laser treatment and account for general variance, we quantified and statistically tested only data that crossed this ± 10% criterion of the normalized pre-measurements (see dashed lines in Fig. 3). In the C1V1 group, VTA stimulation led to a significant increase of peak amplitudes in infragranular layers Vb/VIa most prominently after BF-stimulation for up to 45 minutes. The main current sink in thalamocortical input layers III/IV, in contrast, showed only minor modulation after VTA stimulation. Only 30 minutes after VTA stimulation off-BF evoked activity was increased. Activity in layer Va was unaffected by VTA stimulation. Late onset sink components in layers I/II and VIb showed opposing temporal effects after VTA stimulation. While supragranular activity increased gradually and long-lasting mainly for BF/near-BF stimulation, infragranular layer VIb activity was most strongly amplified for non-BF only during combined tone-laser stimulation. In the YFP-group peak amplitudes of all sink components showed a general trend to be reduced during laser-stimulation and recovered over time within the >45min of post recordings. These effects were potentially due to light-related heating of the stimulation site [25, 26], which might shift the balance of excitation and inhibition in the VTA [27–30]. Data from control animals displayed a mainly stable amplitude range within the ± 10% range of pre-measurements.

**Figure 3 -.**
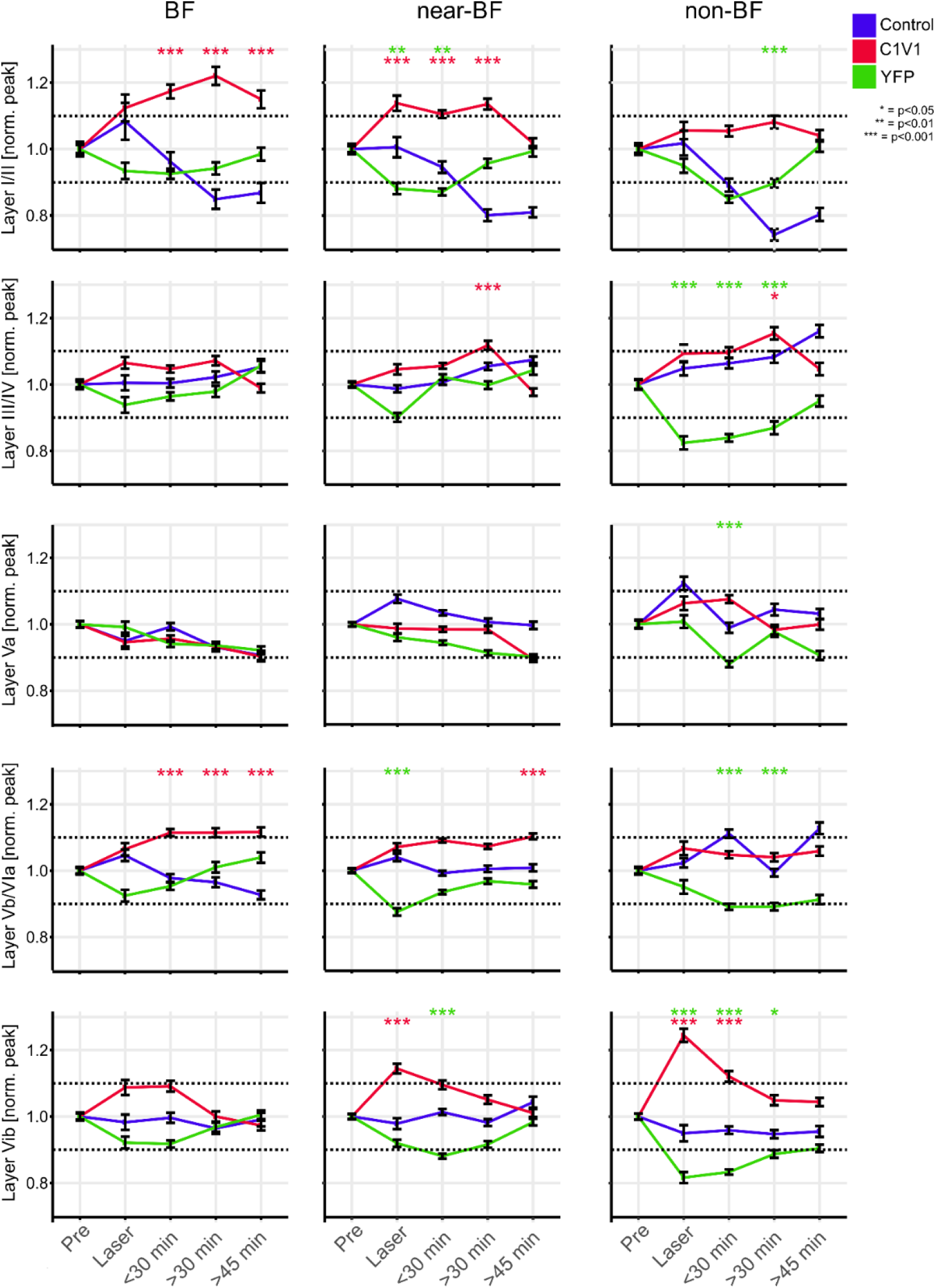
Binned time courses of single-trial sink peak amplitude data of layer-specific CSD-traces. Temporal development of single-trial sink peak amplitudes for early sink activity (sink onset <50 ms) in layers III/IV, Va and Vb/VIa and late sinks (sink onset >50 ms) in layers I/II and VIb for BF, near- and non-BFs. Layer I/II peak amplitudes showed a significant increase for the C1V1 group during laser and in post-measurements, surpassing the +10% criterion within the BF and near-BF bins. Significant differences from the control group are again revealed by LME models and indicated by asterisks. Layer I/II peak amplitudes in the YFP and control group did not show corresponding increases over the recording time but showed a decrease and recovery for the YFP group. Layer III/IV activity did not reveal differences over time for BF-evoked responses in all groups. With spectral distance from the BF the C1V1 group showed a moderate increase yielding significance at the >30 min time bin compared to control animals. Within the non-BF bin, peak amplitudes showed a significant decrease in the YFP group during and after laser stimulation. Layer Va peak amplitudes were most stable across time and groups only displaying a significant decrease for the YFP group in the non-BF bin at <30 min. Layer Vb/VIa peak amplitudes in the C1V1 group were significantly increased in all post-measurements after BF-stimulation. For stimulation with near-BF a similar trend was found. Significant changes between the YFP and control group were again due to decreased peak amplitudes in the YFP group. Layer VIb displayed a highly significant increase in the C1V1 group during laser stimulation, most prominent in the non-BF bin, and less prominent in near-BF and BF bins. Laser-induced increase only persisted for the next time bin, but then recovered to pre-condition. While the control group showed stable peak amplitudes over the time course, YFP animals again showed a moderate decrease most prominent in off-BF bins.

Based on the most prominent changes of evoked activity in the C1V1-group during VTA stimulation (layer VIb) and over the longer period of >30 min after stimulation (layers I/II and Vb/VIa), we compared slopes of evoked peak amplitudes as a function of stimulation frequency (Fig. 4). All data were normalized to BF-evoked amplitudes of pre-measurements within each layer separately. Slopes across the BF/near-BF/non-BF range therefore allow us to characterize changes in tuning sharpness. Frequency-independent gain increase, in contrast, would not affect the slope, but rather affect the intercept. In the pre-measurement, the most prominent frequency tuning was observed in thalamocortical input layers III/IV. In the pre-condition amplitudes in the non-BF bin were reduced about 60% compared to BF-evoked responses. Across frequency bins, this led to a slope of 0.28 indicating a significant dependence of amplitude and stimulation frequency (Fig. 4, blue line, p<0.001). During laser stimulation and 30 minutes after, slopes did not change (change of slope >0.01; p>0.05) indicative of a stable frequency tuning in layers III/IV. In layers Vb/VIa, a slope of 0.13 indicated a less prominent frequency tuning in the pre-condition. During VTA stimulation the frequency tuning was unaffected (change of slope <0.01; p<0.05) but showed a significant increase of sharpness 30 minutes later (change of slope pre:30 min of +0.05; p<0.001). Comparably, in supragranular layers, I/II slopes of the pre-condition (0.09; p<0.001) did not significantly increase during VTA stimulation (change of slope +0.04; p=0.09), but significantly increased >30 min later (change of slope pre:30 min +0.08, p<0.001). Hence, cortical layers Vb/VIa and I/II showed no change of tuning bandwidth during the paired tone/VTA stimulation, but a significant increase in spectral integration prolonged for up to 30 minutes afterwards. Activity in layer VIb showed a prominent increase during VTA stimulation selectively for the off-BF bins (cf. Fig. 3; bottom right), and, in accordance, we found slopes to be significantly decreased during laser stimulation (change of slope pre:laser −0.06; p<0.001), which recovered after 30 minutes (change of slope Pre:30 min −0.02, p=0.10). Layer Va similarly showed a decrease in slope during laser-stimulation (change of slope pre:laser −0.04; p<0.001), and also recovered 30 minutes later (p>0.05). VTA stimulation yielded a transient increase in response bandwidth in cortical layers Va and VIb.

**Figure 4 -.**
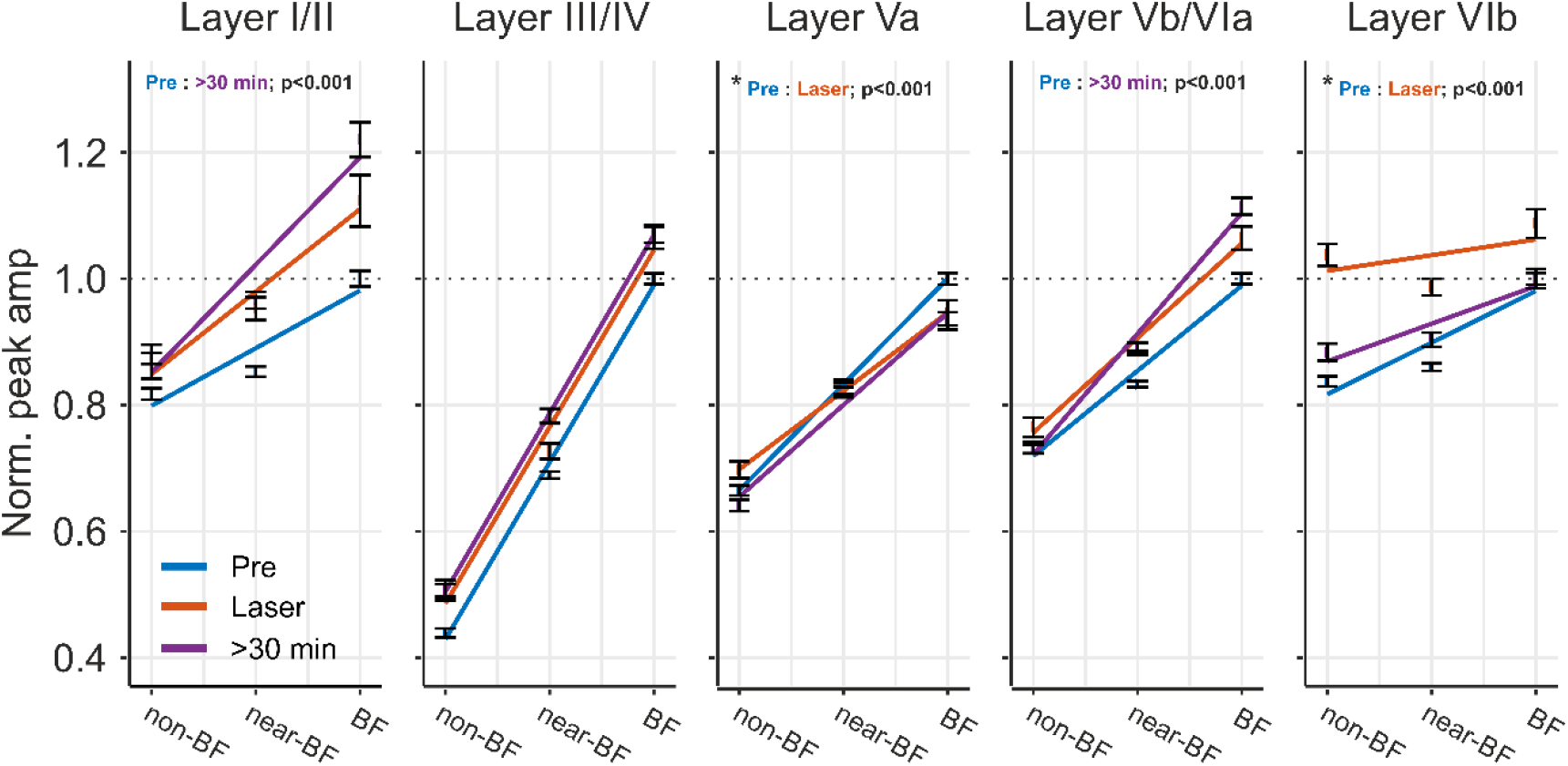
Layer-specific early and late changes of tuning sharpness of the C1V1 group. For data from the C1V1 group, peak amplitudes within each layer were normalized to BF-responses before laser stimulation. We fitted a linear line across evoked responses across the BF-, near-BF- and non-BF-bin and calculated the slope as indicator for spectral tuning sharpness (for pre, laser, >30 min). Sharpest tuning properties of peak amplitudes were found in layers III/IV (slope of 0.28). Tuning in layers III/IV was not affected by VTA stimulation. Layers I/II showed less prominent tuning revealed by a shallower slope in the pre-condition. While VTA stimulation did not yield an immediate significant difference, slopes indicated a significantly sharper tuning >30 min after VTA stimulation (p< 0.01). This was due to mainly an increase in BF-evoked peak amplitudes in accordance with Fig. 5. Peak amplitudes in infragranular layers Vb/VIa showed a similar effect as layers I/II with a trend of sharper tuning during VTA stimulation that yielded significance >30 min later. In contrast, only in late infragranular layer VIb, peak amplitudes revealed a significantly shallower slope indicative of a broader frequency tuning during laser stimulation (Pre:laser; p<0.01), which recovered after >30 min (Pre:<30 min; p=0.1). This can be seen by the flat slope between non- and BF bins. Layer Va only showed a minor decrease of the slope during VTA stimulation that recovered partially >30 min later. Significances indicated are based on slope comparisons performed using the Ismeans package in R (version 2.27-62). For statistical explanation, see text.

Based on the layer-specific modulation of early tone-evoked inputs in thalamocortical-recipient layers Vb/VIa and subsequent synaptic inputs in supragranular layers, we speculate that stimulation of the VTA affects early and late sensory processing on a local and corticocortical level, respectively [cf. 13].

### Impact on local intracolumnar and corticocortical synaptic inputs

In a next step, we therefore aimed to disentangle early activity associated with local thalamocortical input, and subsequent activity associated with corticocortical processing. We compared changes in the tone-evoked average rectified CSD (AVREC) and the residuals of the CSD (ResidualCSD) during early (0-50 ms) and late (80-300 ms) time windows (Fig. 5 A). This rationale is based on previous findings demonstrating that the residual CSD reflects the spatiotemporal ratio of unbalanced sinks and sources providing a quantitative measure of horizontal intercolumnar inputs to a given cortical site [15, 21]. The AVREC, on the other hand, reflects the overall intracolumnar current flow at a given recording site [31, 32]. Fig. 5 A (left) shows changes of both the grand average of the AVREC (top) and relative CSD (bottom) waveforms before, during, and, in certain time windows, after VTA stimulation in the C1V1 group. Comparable changes were not observed in neither the YFP nor control group (Additional file 3). To further investigate temporal changes of BF-evoked responses in more detail, time course of binned root mean square (RMS) values for early and late averaged and normalized AVREC and ResidualCSD windows of each group were plotted as a function of time (normalized to pre-measurements). RMS values of the AVREC during the early phase (0-50 ms) displayed a significant increase in the C1V1 group from 15 min until 52 min after VTA stimulation. Significant effects were validated with a LME model (Fig. 5 B; p<0.001-0.025). During the late phase (80-300 ms) after tone onset, RMS of the averaged AVREC did not show any significant changes between each group (Fig. 5 B). During laser stimulation, a significant increase has only been observed for the averaged early ResidualCSD indicative of amplified lateral contributions to early tone-evoked columnar activity in the C1V1 group (p=0.04). Furthermore, early and late ResidualCSD were both significantly enhanced during later time points after the VTA stimulation (early ResidualCSD: 22.5-37.5 min; p=0.014-0.039; late ResidualCSD: 37.5-45 min, p=0.008-0.033). Averaged RMS tuning curves of the AVREC and ResidualCSD of the C1V1 group revealed an increase of tone-evoked activity during and after laser stimulation across the entire range of BF, near-BF (± 1-2 octaves) and non-BF (± 3-4 octaves) stimulation (Fig. 5 C).

**Figure 5 -.**
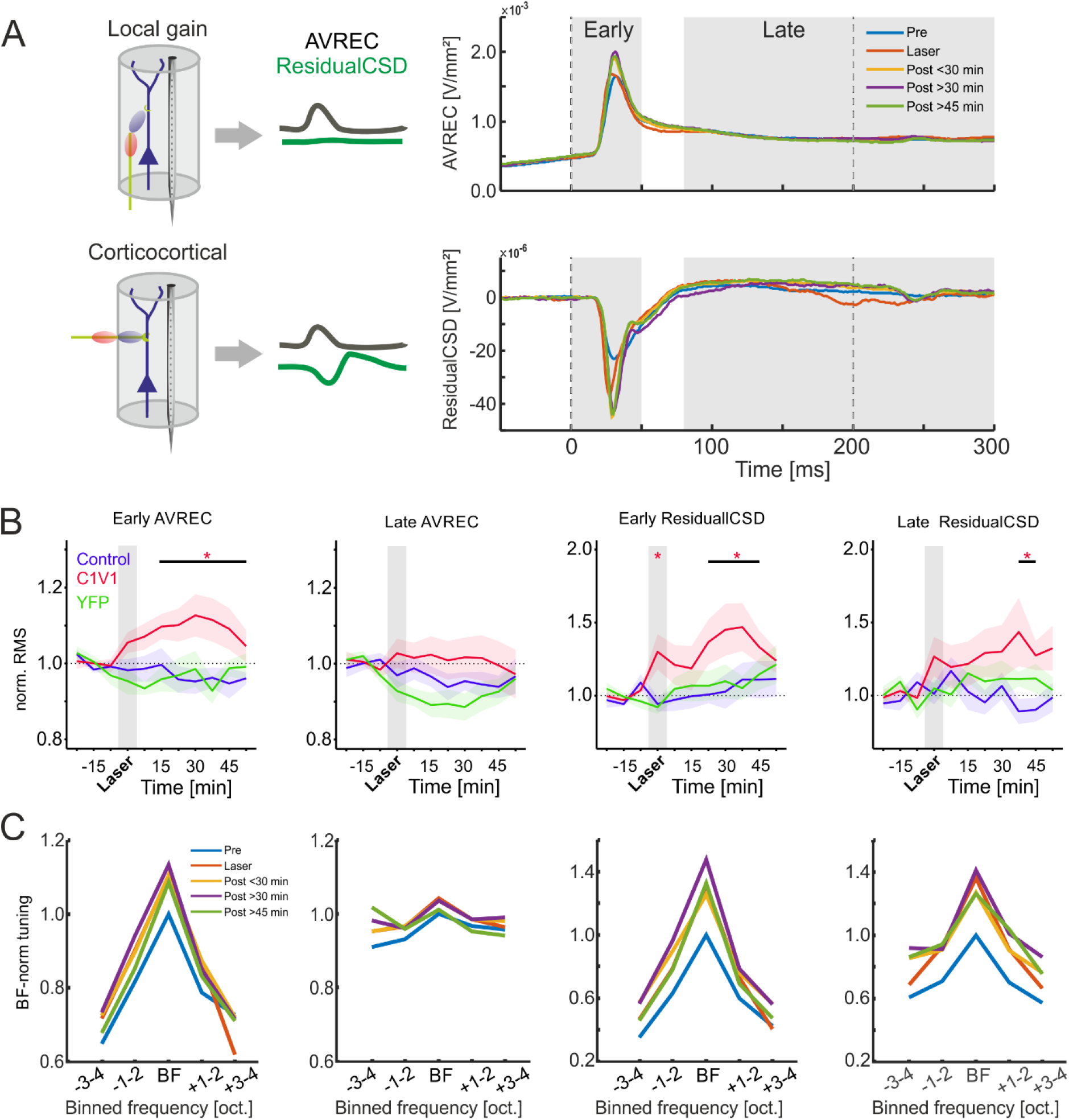
Temporal development of early and late AVREC and ResidualCSD activity. (A) Grand average traces of the BF-evoked AVREC (top) and ResidualCSD (bottom) of the C1V1 group (n=12) for binned pre, laser and post (<30, >30 &>45 min)-measurements. Right, insets illustrate that AVREC activity is mainly associated with local columnar activity, while the ResidualCSD quantifies contributions from lateral corticocortical input (cf. Happel et al., 2014, 2010). (B) Temporal time courses of Pre-normalized early/late AVREC and early/late ResidualCSD traces of C1V1 (red), control (blue) and YFP (green) groups (n=12, n=7, n=7, respectively) shown for BF stimulation. Early AVREC signal displayed a significant increase in post-measurements for the C1V1 group whereas time courses of the control and YFP groups were stable. Late AVREC signal showed no significant differences between either group. RMS ResidualCSD values of early and late time windows displayed a similar temporal development as the early AVREC becoming significantly increased in the later post-measurements within the C1V1 group. Control and YFP groups showed no significant changes over time. Note the peaking in the early ResidualCSD RMS during laser stimulation, which is only present in the C1V1 group. Asterisks mark significant changes according to the applied mixed linear models of the normalized data towards the control group (see Materials and Methods). (C) Tuning curves of the RMS value normalized to BF-evoked responses in the pre-condition for early/late AVREC and early/late ResidualCSD for binned time points for C1V1 group. Note the general increase of the overall tuning curve relative to pre measurements.

### Integration of local and corticocortical synaptic inputs in dependence of the stimulation frequency

All single-trial AVREC and ResidualCSD RMS values were normalized to pre measurements. Normalized values were analyzed as a function of time before and after VTA stimulation binned for BF, near-, and non-BF stimulation for all groups (Fig. 6). Fig. 6 A (top) shows a gradual increase of the early AVREC compared to the control- and YFP-group independent of stimulation frequency. For BF- and near-BF-stimulation, this yielded a significant increase at >30 min after VTA stimulation. The ResidualCSD also showed a strong increase almost independent of the stimulation frequency already during the combined tone-laser presentation and all following post measurements. Consistently with the layer-specific activity, there was a less pronounced decrease of the near/non-BF-evoked AVREC in early and late time windows in the YFP-group during laser stimulation, which started to recover over time (Fig. 6 A, bottom).

**Figure 6 -.**
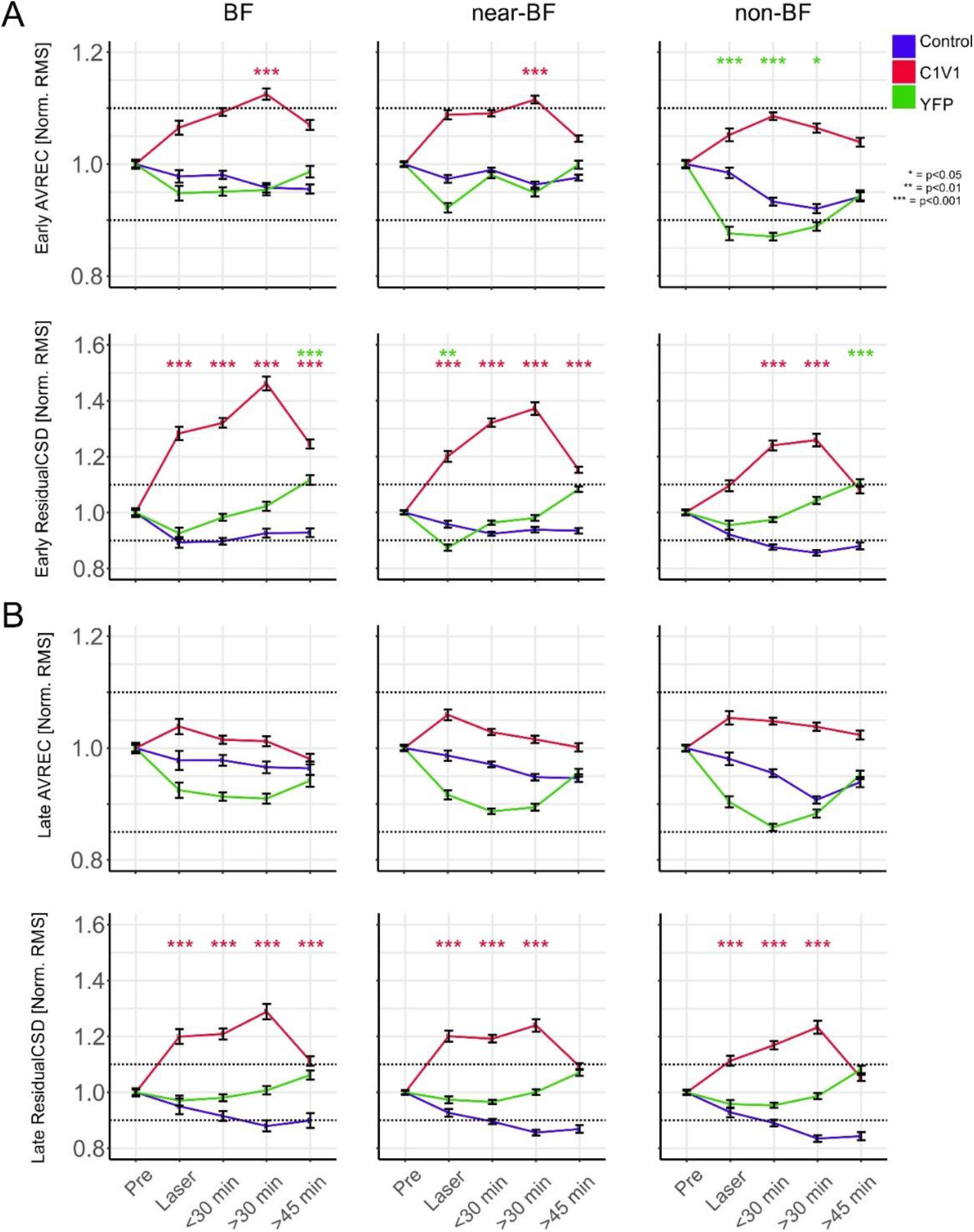
Normalized single-trial early and late AVREC and ResidualCSD RMS values over time. (A) Temporal time course of averaged early single-trial RMS AVREC (top) and ResidualCSD (bottom) values normalized to pre-measurements. Data for BF, near-BF and non-BF stimulation is shown separately and was binned for pre-measurements, laser, and post-measurements (<30 min, >30, >45 min). Note the prominent increase of early RMS AVREC values for the C1V1 group whereas control and YFP group displayed a more stable or decreasing time course, respectively. Also, early RMS ResidualCSD values significantly increased during and after the Laser measurements for BF, near- and non-BF bins of the C1V1 group, whereas YFP and control display a more stable time course. (B) Temporal time course of late Pre-normalized RMS AVREC and ResidualCSD values of single-trial data for BF, near-BF and non-BF. Late RMS AVREC values for the C1V1showed a mild, yet not significant, increase that did not surpass the +10% criterion. The control group displayed a stable and slightly declining time course for the late RMS AVREC values, while the YFP group showed a moderate decrease that recovered over time. Late RMS ResidualCSD values increased during and after the Laser measurements for BF, near- and non-BF bins of the C1V1 group, whereas YFP and control groups both showed a more stable time course. Significances in A) and B) were calculated using mixed linear models on the normalized data. Asterisks indicate corresponding significances in comparison to the control group, in case values exceed the ± 10% pre-criterion. Single-trial CSD data in following figure is presented equally.

In the later time window of 80-300 ms after tone-onset, only the ResidualCSD showed a significant increase during and up to 30 minutes after VTA stimulation (Fig. 6 B). Altogether, VTA stimulation led to a gradual increase of the overall early current flow most pronounced for BF- and near-BF stimulation. This effect was long-lasting and peaked around 30 minutes after stimulation. We further found an immediate and long-lasting increase of imbalance between sinks and sources based on the early and late ResidualCSD in all frequency bins indicative of amplified corticocortical activity.

## Discussion

The VTA is a key structure to convey information about stimulus salience and valence to target areas distributed throughout the brain [2]. Receiving both sensory as well as VTA inputs, the primary sensory cortex is ideally positioned to integrate such bottom-up and top-down information to shape behavior through reinforcement learning. We here demonstrate for the first time that activation of VTA-reward circuits amplifies sensory input via a yet largely overlooked thalamocortical projection targeting infragranular layers Vb/VIa of the auditory cortex [22, 33]. In contrast, granular layer III/IV input, a well-known substrate for sensory gain modulation [14, 15, 33, 34], was unaffected by VTA stimulation. The impact of the observed infragranular gain modulation on supragranular layers manifests as a translaminar long-lasting enhancement of frequency tuning (see Fig. 7). Furthermore, we demonstrate that the frequency-specific gain in deeper layers emerges on a local columnar level and yields an increased corticocortical integration displayed in lateral input most likely terminating in upper layers. This highlights the diverse, and yet elusive functions of cortical layers in the sensory cortex [35]. The long-lasting modulation after VTA stimulation reveals that the influence on sensory cortex transcends the time scales commonly associated with reward-prediction error coding.

**Figure 7 -.**
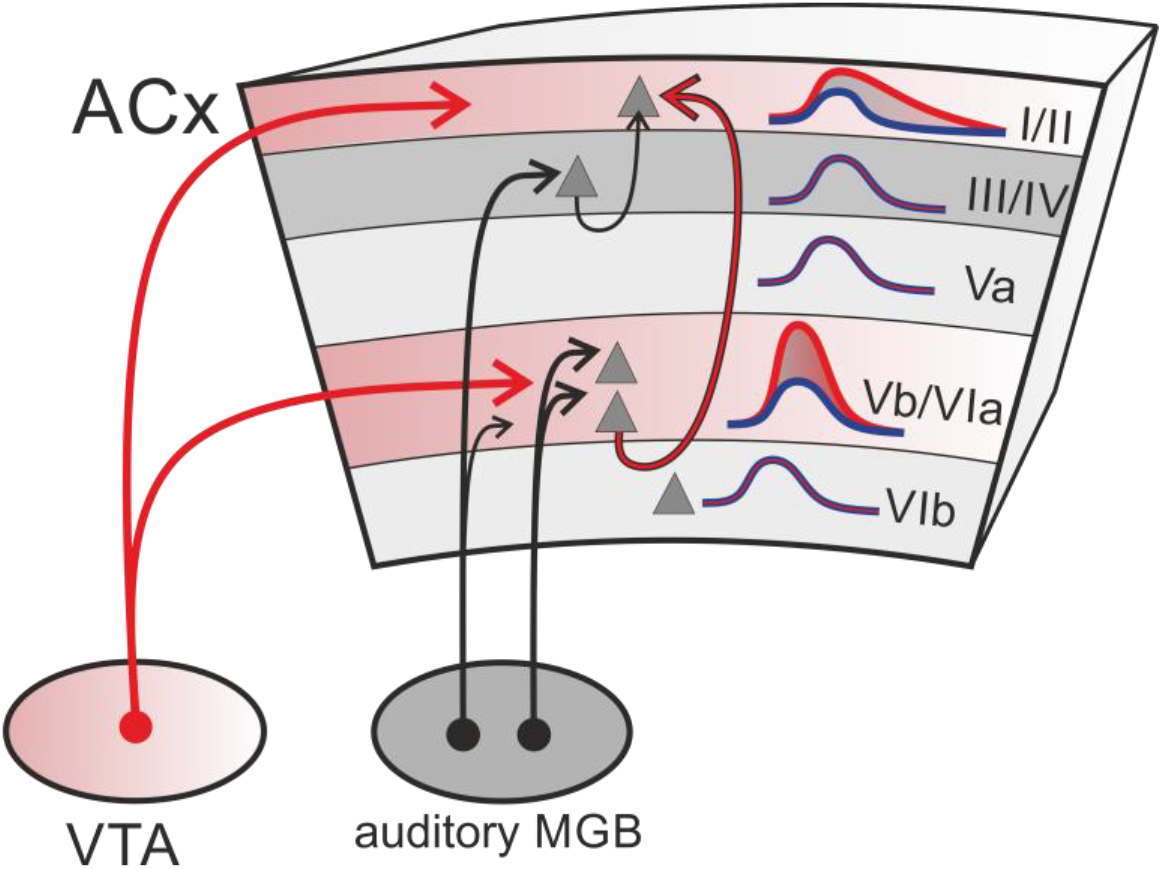
Schematic illustration of convergent deep layer inputs from sensory thalamus and VTA. VTA projections mainly target upper and infragranular layers of the gerbil auditory cortex (see also Fig. 1 B). Stimulating the VTA projections led to a frequency-specific gain amplification of early sensory-evoked responses in thalamocortical recipient layers Vb/VIa. This gain increase effectuated an enhanced spectral tuning representation in supragranular layers I/II. The supragranular gain modulation might hence be inherited via intratelencephalic neurons targeting upper layers and bypassing granular input circuits [36], may originate from a direct influence of dopamine released in upper layers or a combination of both. MGB, medial geniculate body.

### VTA-activation enhances infragranular thalamocortical inputs in the auditory cortex

Optogenetic stimulation of VTA-projection neurons allowed us to investigate effects of reward-related cross-regional circuits (Fig. 1) [19]. Roughly 8% of non-sensory inputs of gerbil ACx arise from direct projections of the ipsilateral VTA, which terminate across the entire cortical column [4]. Therefore, direct stimulation of the VTA projection neurons is the most effective method to activate this circuit. In contrast, the success of optogenetic approaches to directly stimulate cortical terminals have been challenged in recent studies and particularly do not allow for an homogenous activation across all cortical layers [26]. VTA stimulation led to increased early tone-evoked (0-50 ms) local columnar current flow and increased long-range corticocortical activity. This effect persisted for up to 50 minutes after VTA stimulation and was most prominent for the respective BF of the given recording site (Fig. 5). Hence, VTA stimulation promotes the local gain of columnar input over a prolonged period and lead to substantial corticocortical integration of spectral information. Notably, the gain amplification of sensory input appeared exclusively in thalamocortical-recipient layers Vb/VIa but was absent in the main thalamocortical-recipient granular layers III/IV (Fig. 3). This enhancement of infragranular activity was locally relayed to supragranular layers I/II. This mechanism may provide a way for intratelencephalic neurons in infragranular layers [36] to provide a translaminar gain control of sensory representation depending on the experience of reward [37, 38]. Although direct projections of the ipsilateral VTA provide a significant contribution to non-sensory inputs of the gerbil ACx [4], our approach may recruit other relay stations that could contribute to the observed effects.

Previously, reported effects of dopamine on tuning sharpness in the ACx [39, 40] raised the question on which specific cortical circuit motifs reward information modulates spectral information. Our data showed that stimulating the VTA projections only affected early auditory tuning sharpness in deep cortical layers, but not in the main thalamocortical input layer III/IV [33, 41]. Sharper frequency representation was relayed to superficial layers I/II and persisted over time after VTA stimulation. Our observations are thereby consistent with recent findings from layer-specific high-field fMRI in human auditory cortex showing sharper tuning in upper layers depending on attention and task engagement [42].

As a result, reward-circuit activation promotes a long-lasting gain enhancement of salient auditory stimuli via strengthened infragranular thalamocortical inputs and increased sensory integration in supragranular circuits. Our finding now provides an explanatory role of the anatomically and physiologically well-described early infragranular inputs in layers Vb/VIa of the sensory cortex [15, 22, 33, 43] as a circuit for integrating sensory and task-related information.

### Persistent cortical modulation might underlie dopamine-dependent influences on learning

Recent reports have highlighted that sensory-evoked responses in deep layers are modulated by behavioral states [44, 45] and undergo substantial learning-related plasticity [46] not described for middle layer thalamocortical input [47]. Our data now assigns a functional impact of the VTA on deep cortical layers—particularly under the control of dopaminergic neuromodulation (Fig. 7).

Optogenetic VTA stimulation preferentially recruits the main output population of dopaminergic projection neurons making up roughly 55-65% of all VTA neurons [48–50]. Nevertheless, it has been demonstrated that the small population of non-dopaminergic projection neurons also convey reward information [51–53] and particularly promote the activation of brain-wide regions including the auditory cortex [19]. Furthermore, there is overlap between dopaminergic and glutamatergic cells through transmitter co-release [54–56]. Nevertheless, the long duration of responses modulation in our data set strongly argues for a significant dopaminergic component, as glutamatergic effects are generally more transient [57]. Further evidence for a dopaminergic origin of the long-lasting effects is provided by the distribution of dopamine receptors. In the sensory cortex, they are concentrated mainly in infragranular and supragranular layers where we see modulatory effects (Fig. 1 B) [11, 58–62].

## Conclusions

Phasic release of dopamine from the ventral tegmental area has long been associated with the reward prediction error signal postulated by reinforcement learning theory. How dopamine released by the VTA affects the plasticity that integrates sensory information and such learning-related signals in sensory cortex, however, is unknown. Our findings therefore assign a specific function to this rather understudied cortical circuit: VTA-modulated input to deep layer neurons in the sensory cortex mediates the integration of sensory and task-related information even when they are not precisely temporally contingent. Our study pursues the idea of a VTA-mediated local gain of sensory input at infragranular thalamocortical terminals via presumably presynaptic dopamine receptors [13, 63, 64]. Previous studies have described accumulative levels of dopamine in the auditory cortex over 30 minutes after training onset impacting on learning and long-term memory consolidation [9] suggesting a long-lasting action of dopamine at the presynaptic terminals [11]. Thereby, the sensory gain amplification within deep layer neurons in the sensory cortex described in this study potentially promotes a sustained perceptual salience of incoming stimuli [65–67]. Our findings now provide a circuit level explanation of the necessary integration of sensory and task-related information even when they are not precisely temporally contingent. Thereby, this circuit might constitute a substrate for the credit assignment in reinforcement learning theory concerned with the temporal delay between sensory input and its behavioral consequences [68].

## Methods

All experiments were carried out in adult male Mongolian gerbils (*Meriones unguiculatus*, age 4-8 months, body weight: 70-100 g; n= 26). The C1V1 group (n=12) consisted of animals that were expressing the viral construct, allowing for optogenetic stimulation of the VTA whilst recording. The control group (n=7) consisted of non-transfected animals to control for temporal stability of the signal. In order to investigate effects due to laser-stimulation itself, an opsin-less YFP group (n=7) was tested. Experiments were conducted in accordance with ethical animal research standards defined by the German Law and approved by an ethics committee of the State of Saxony-Anhalt.

### Viral transfection of the VTA

Gerbils were anesthetized by intraperitoneal injection (0.06-0.08 ml/h) of 45% v/v ketamine (50 mg/ml, Ratiopharm GmbH), 5% v/v xylazine (Rompun, 2%, Bayer Vital GmbH) and 50% v/v of isotonic sodium-chloride solution (154 mmol/l, B. Braun AG). Animals were fixed in an automated stereotaxic injector (Neurostar) for virus transduction (700 nl; 100 nl/min; 5 min resting; C1V1: AAV5-CamKIIα-C1V1(E162T)-p2A-eYFP or YFP: AAV2-CamKIIα-eYFP 3E12 particles/ml, UNC Vector Core; target side: AP: −3.8-4.0; ML: 0.5; DV: −6.2 mm, bregma for reference) and fiber implantation with a planar setting of bregma and lambda. Custom made fiber implants (fiber implant length 8 mm, Ø 230 μm, NA 0.39) were implanted at target side (AP: - 3.8-4.0; ML: 0.5; DV: −5.8 mm). Fiber positions were determined by histological analysis and compared with the gerbil brain atlas (Fig. 1 D and E) [69]. Animals were kept for 3 weeks to allow recovery and enough protein expression.

### Intracranial self-stimulation using optogenetic VTA activation

Successful activation of the VTA through optogenetic stimulation was confirmed in an intracranial self-stimulation paradigm. Animals were trained for 10 consecutive days (at 20 min per day). By activating a lever with their nose, they could elicit brief laser stimulation (10 pulses, 25 Hz, 473 nm, 10 mW) delivered to the VTA through the implanted fiber (Fig. 1 A, B and C). Only animals pressing the lever more than 50 presses/minute were kept for the C1V1 group (Fig. 1 G) [18–20].

### Electrophysiological recordings

Surgical procedure for electrophysiological recordings have been described in detail elsewhere [40]. Briefly, animals have been anesthetized by intraperitoneal injection (0.06-0.08 ml/h) of 45% v/v ketamine (50 mg/ml, Ratiopharm GmbH), 5% v/v xylazine (Rompun, 2%, Bayer Vital GmbH) and 50% v/v of isotonic sodium-chloride solution (154 mmol/l, B. Braun AG). The right auditory cortex was exposed by trepanation and target location of AI was chosen based on vascular landmarks. Status of anesthesia was checked via paw withdrawal-reflex on a regular basis (10-20 min). Body temperature was kept stable at 34°C. Due to an epileptic seizure during the electrophysiological recordings we had to omit an animal from the YFP group.

Experiments have been carried out in a Faraday-shielded acoustic soundproof chamber. The animals head was fixed 1m away from the speaker (Tannoy arena satellite KI-8710-32) by screwing the head post to a custom-made head holder. Local field potentials (LFPs) have been recorded with a linear 32-channel shaft electrode (NeuroNexus A1×32-50-413), which was implanted perpendicular to the surface area of AI [15]. LFPs have been pre-amplified 500-fold and band-pass filtered (0.7-300 Hz) with a PBX2 preamplifier (Plexon Inc.). Data were digitized at a sampling frequency of 1 kHz with a multichannel recording system (Multichannel Acquisition Processor, Plexon Inc.).

Pure tones spanning 8 octaves (frequency range between 125 Hz-32 kHz, tone duration: 200 ms, ISI: 0.6-0.8, 50 pseudorandomized repetitions, 65 dB SPL, 7.5 min per measurement) were generated in MATLAB, converted into an analog signal by a data acquisition card (NI PCI-BNC2110, National Instruments, Germany), rooted through a controllable attenuator (gPAH, Guger, Technologies, Austria) and amplified by an audio amplifier (Thomas Tech Amp75). A measurement microphone and conditioning amplifier were used to calibrate acoustic stimuli (G.R.A.S. 26AM and B&K Nexus 2690-A, Bruel & Kjaer, Germany).

Electrophysiological measurements were performed according to the scheme of Fig. 2 A. After implantation, laminar LFPs were measured for 45-75 min until evoked CSD profiles had been stabilized (Pre-measurements). The last three recordings before laser treatment were taken as the referencing Pre-measurements. For the C1V1and YFP groups, light fibers were additionally connected to a laser for VTA stimulation. Laser stimulation (10 mW at the optical fiber of the setup) was then coupled and synchronized with tone presentation for one measurement (25 Hz, 473 nm, 10 mW; 400 repetitions in total). This measurement was taken as the laser measurement followed by a series of post-measurements (>60 min). After the experiment, gerbils were perfused with 4% PFA as described previously [70] and brains were removed for anatomical and histological validation.

### Current source density analysis

Based on the tone-evoked LFPs for each presented stimulus we calculated the second spatial derivative, indicating the laminal evoked current-source density (CSD) distribution in AI [71]:

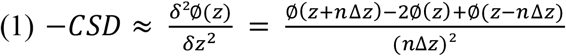

The formula reflects the relation of the field potential (*Ø*), the spatial coordinate of the cortical laminae (*z*), the inter-channel distance (Δ*z*, 50 μm) and the differentiation grid (*n*). Prior to CSD calculation, LFPs were smoothed using a weighted average of 9 channels (Hamming window, spatial filter kernel size of 400 μm; linear extrapolation of 3 channels at boundaries) [15].

CSD profiles revealed patterns of current influx (sinks) and efflux (sources). Based on CSD profiles of the Pre-measurements, cortical layers were assigned to the respective channels of sink components. Early sink components were used as indicators of layers III/IV, Va, and Vb/VIa, whereas later sink activities were assigned to layers I/II, and VIb (Fig. 2; Additional file 2).

CSD data was also transformed by rectifying and averaging the waveforms of each single channel (*n*). This average rectified CSD (AVREC) provides a temporal waveform of the overall local columnar transmembrane current flow [31, 32].

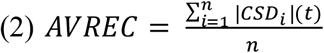

We further quantified the residue of the CSD defined as the sum of the non-rectified magnitudes divided by the rectified magnitudes for each channel. Thereby, the ResidualCSD quantifies the balance of the transmembrane charge transfer along the recording axis [72] and gives rise to the lateral corticocortical contribution to stimulus related activity [15].

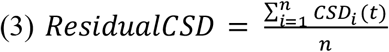

In order to quantify the early (0-50 ms) and late (80-300 ms) contributions of the AVREC and ResidualCSD, root mean square (RMS) amplitudes were calculated. Based on the previously described layer indexing, averaged layer-dependent CSDs were calculated, which were used for the semi-automatic sink detection (Additional file 2). Briefly, before detection of significant tone-evoked activity, sources were omitted before calculating the mean of CSD traces within a cortical layer. Then, the 1.5-fold standard deviation (SD) of the baseline (−200-0 ms before tone presentation onset for all channels and stimuli) was used to calculate the intercepts of the smoothed (10 ms gaussian sliding window) mean CSD trace. For early tone-evoked sink activity in cortical layers III/IV, Va, and Vb/VIa, we used an onset of the first intercept between 0-50 ms after tone onset. For subsequent tone-evoked sink components in layers I/II and VIb onsets were only taken >50 ms after tone-onset. Offsets of each sink were defined as the crossings below the 1.5 SD baseline and for each cortical layer. Peak amplitudes of layer-dependent CSD-traces were determined within time windows of on- and offsets from the average of the corresponding CSD channels. For calculating the RMS value of AVREC and ResidualCSD we used fixed time windows of 0-50 ms after tone-onset for early and 80-300 ms for late activity.

### Data sorting and analysis

Tuning curves of layer-specific CSD-traces, AVREC, and ResidualCSD were averaged according to the best-frequency of each animal separately to plot an averaged group tuning curve. For illustration, all non-BF frequencies above and below the BF were binned pairwise, resulting into the grouping of near-BFs (± 1-2 octaves) and non-BFs (± 3-4 octaves) in 5 frequency bins [− non-BF, − near-BF, BF, + near-BF, + non-BF]. For further analysis of BF-, near- and non-BF evoked responses, we observed data from time windows before, during, and after laser stimulation. Evoked responses were normalized within each animal using the corresponding mean values of the last three Pre-measurements before laser-stimulation (Fig. 2 A.

For statistical comparison, parameters of interest were analyzed on a single-trial level using linear-mixed effect (LME) models (R Studio; R 3.5.1) [24] binned across measurements as follows: pre 1-3 (−22.5 - −7.5 min), laser (0 min), post 1-3 (7.5-22.5 min), post 4-5 (30-37.5 min), post 6-7 (45-52.5 min). Peak amplitudes of single-trial data were determined within the time windows from on- and offsets revealed on averaged CSD traces. LME models were calculated and fitted using normalized data with the lmerTest-package (3.0-1) and a Wald-chi-square-test was performed using the car-package (3.0-2). Animals were used as “by-subject” random effects, to account for the repeated measurements during and across time points whereas measurements and groups were used as fixed effects to explain physiological data. As significance criterion for effects on single-trial level, the normalized signal needed to i) exceeded a ± 10% criterion compared to Pre-measurement (based on the SEM of the average jitter before laser treatment), and ii) show significant difference from control group data revealed by significant fixed effects of the LME models. For slope estimations and comparison between measurements, we used the R-package Ismeans (version 2.27-62) to calculate a least square linear fit for non-, near- and BF data bins and check significant changes in tuning slope properties between measurements.

### Anatomical estimation of fiber position

PFA fixated brains of the C1V1 group (n =7) were cut in 50 μm thick slices around the region of the VTA. Every second slice was used for Nissl staining to anatomically evaluate the fiber position, whereas the remaining slices were used for immunostaining to determine the distance from fiber tip to areas of tyrosine hydroxylase (TH) and YFP-co-fluorescence with FIJI ImageJ (1.51g). Pictures were taken with a Leica microscope (Axioscop 2, Leica GmbH).

Immunostaining was performed as previously described [19] with a primary antibody against tyrosine hydroxylase (1:1000 rabbit anti-TH, Milipore) and a secondary Alexa Fluor 546 goat-anti-rabbit (1:400 anti-rabbit, Molecular probes) using an antibody diluent (Zytomed Systems). Position of the fiber was estimated using the recently published gerbil brain atlas for reference [69].

## List of abbreviations

ACx: Auditory cortex
AI: Primary auditory cortex
AVREC: Average rectified CSD
BF: Best frequency
C1V1: Animal group expressing a functional chimeric opsin in the VTA
DA: Dopamine
CSD: Current source density
LFP: Local field potential
LME model: Linear mixed effect model
PFA: Paraformaldehyde
RMS: Root mean square
SEM: Standard error of mean
TH: Tyrosine hydroxylase
VTA: Ventral tegmental area
YFP: Animal group without a functional opsin only expressing a yellow fluorescent protein in the VTA

## Declarations

### Author contributions

FWO, MTL and MFKH designed research. MGKB and KED performed electrophysiology. MGKB and SV performed histology and behavioral experiments. MGKB, KED, MK, MD and MFKH analyzed data. MGKB, KED and MD contributed analytic tools. MGKB and MFKH prepared the figures and wrote the initial draft. MGKB, KED, FWO, MTL and MFKH wrote the manuscript. All authors reviewed and revised the manuscript.

## Acknowledgements

We like to thank Janet Stallmann, Anja Gürke and Kathrin Ohl for assistance with surgeries, histology- and immunostaining of brain slices. We would like to thank PD Dr. Eike Budinger and Dr. Julia Henschke for their help and assistance in anatomical questions. Many thanks to our interns for their assistance with anatomical reconstructions: Tarik Drewes, Katharina Eick, Vincent Koop, Jan-Niklas Löbner, Julia Rieger and Ceylan Steinecke. This project and corresponding research have been funded by the Leibniz Association WGL by the Leibniz Postdoctoral Network, LPN (to MTL and MFKH) and, the Deutsche Forschungsgemeinschaft DFG by the SFB779 (to MFKH and FWO) and SPP1665 (to MTL and FWO).

## Additional files

**Additional file 1-.**
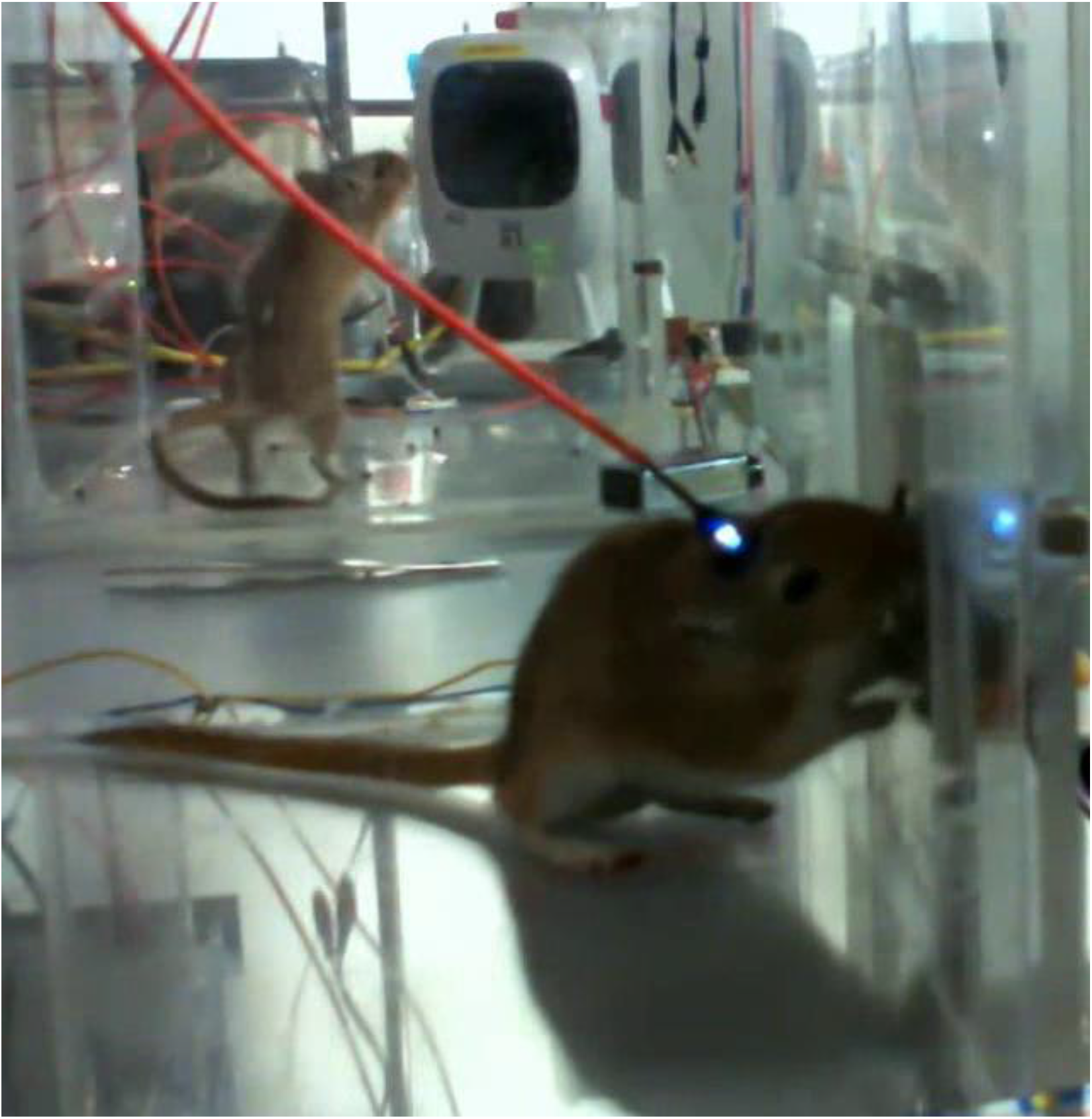
Optogenetic intracranial self-stimulation induced by lever pressing. Animal video clip of an intracranial self-stimulation training session. Animals were placed into a 25×35×22cm^3^ Plexiglas box for 20 min once a day. Optogenetic implants were attached via an optical fiber with a shutter-controlled laser (10 mW, 473 nm). By pressing an installed lever, animals were able to trigger a laser pulse (25 Hz, 5 ms) for VTA stimulation. Highly addicted animals (>= 850 lever presses per session), were kept for optogenetic experiments whereas non-addicted animals were rejected from this study (Compare behavior of the animal in the background).

**Additional file 2-.**
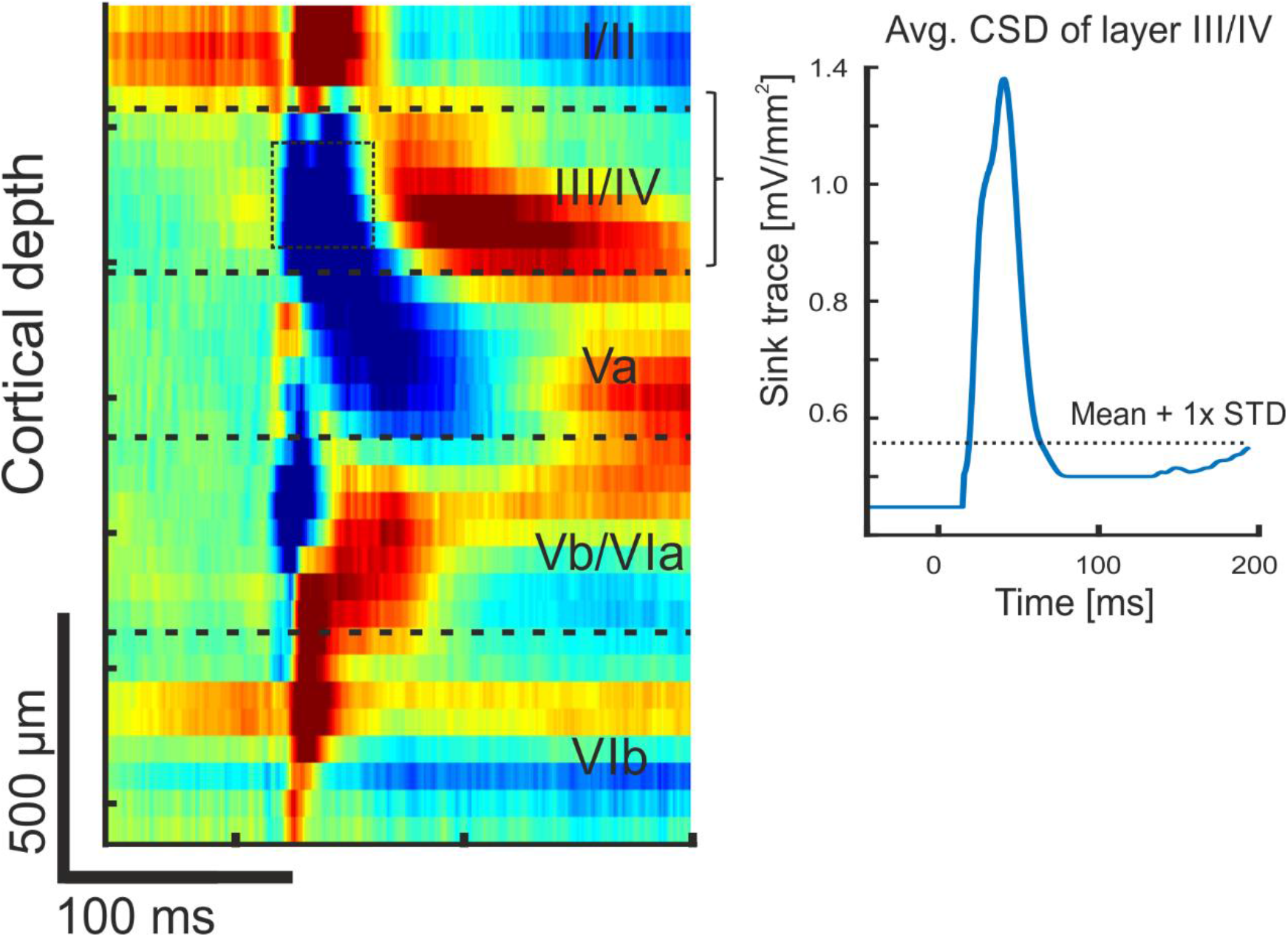
Assignment of layers to CSD channels and semi-automatic detection of sink-activity. Based on previous studies, earliest tone-evoked components are related to layers Vb/VIa and III/IV [15, 22]. Subsequent synaptic activity is conveyed to layer Va and VIb, and layers I/II, as indicated in the representative example. Channels assigned to individual layers were averaged and activity attributed to sources (positive values) were dismissed. For visual purposes, a corresponding average-sink trace from layer III/IV was inverted. Based on the pre-stimulus baseline signals (200 ms), the baseline mean value + 1 standard deviation (STD) was calculated as response threshold for each layer-dependent CSD trace. A deflection that surpassed a 1-fold STD was used to determine the onset and offset of early and late sink activations (see Materials and Methods). Significant sink onset activity within the first 50 ms after tone onset was assigned as early activity, while later onsets were assigned as late activity.

**Additional file 3-.**
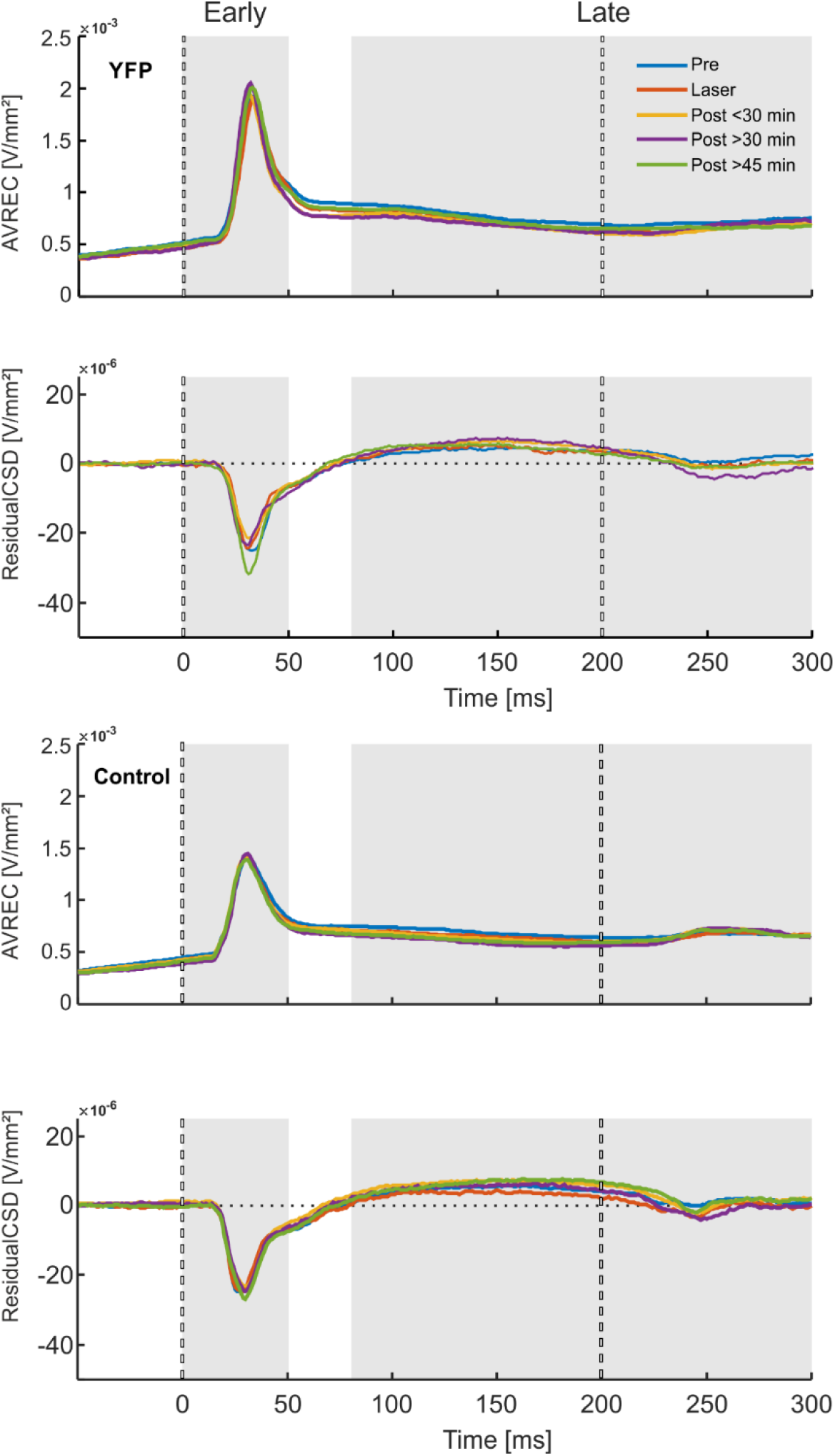
Grand average AVREC and ResidualCSD traces for YFP and control groups. (A) AVREC and ResidualCSD traces of the YFP group (n=7) for binned pre-measurements, laser-measurement, post-measurements < 30 min, post-measurements > 30 min and post-measurements > 45 min. Dashed lines indicate tonal on- and offset. Areas for early and late RMS calculations are underlined with gray. Group sorting was performed according to the BF of the granular sink peak amplitudes. (B) AVREC and ResidualCSD traces of the control group (n=7) for binned pre-measurements, laser-measurement, post-measurements < 30 min, post-measurements > 30 min and post-measurements > 45 min. Dashed lines indicate tonal on- and offset. Areas for early and late RMS calculations are underlined with gray. Group sorting was performed according to the BF of the granular sink peak amplitudes.

